# Latent dentate gyrus engrams formed in infancy scaffold adult learning

**DOI:** 10.64898/2026.10.05.756342

**Authors:** Sam Haar, Cory McKenzie, Sheena A Josselyn, Paul W Frankland

## Abstract

Infantile amnesia refers to the loss of event-based memories acquired early in life^1,2^. In mice, however, these memories are not erased; selective optogenetic stimulation of engrams formed in infancy leads to memory recovery in adulthood^3,4^, indicating that infant-formed engrams can persist in a latent form. We therefore asked whether these latent engrams, although not accessible through natural retrieval, nonetheless influence related adult learning. Using a contextual fear conditioning paradigm, we find that adult learning is facilitated by prior experience in infancy, both for the same and related contexts. Activitydependent engram tagging revealed that adult learning reactivates infant-acquired dentate gyrus ensembles, and selective inactivation of these ensembles during adulthood abolished this facilitation. These findings indicate that memories formed in infancy persist as latent engrams that scaffold subsequent learning, consistent with a role in establishing schema-like representations^5^.

## RESULTS

To characterize infantile amnesia in mice, infant (postnatal day 17; P17) or adult (P56-70) TRAP2 × TdTomato mice were trained in contextual fear conditioning and tested in the same context at either recent (2 days) or remote (28 days) delays (Fig. 1A). Mice received 4-hydroxytamoxifen (4-OHT) immediately after training to permanently label ensembles activated during encoding. Following testing, immunohistochemical analysis of the activity-regulated gene *c-Fos* was used to assess reactivation of these ensembles (TdTomato+/c-Fos+) in the dentate gyrus (Fig. 1B), a key region for this form of learning^6^. Consistent with previous studies^3,4^, mice trained as infants exhibited robust freezing at the recent delay but minimal freezing at the remote delay, reflecting amnesia for these infant-acquired memories. In contrast, mice trained as adults exhibited comparable freezing at both delays, reflecting stable memory (Fig. 1C).

**Figure 1.**
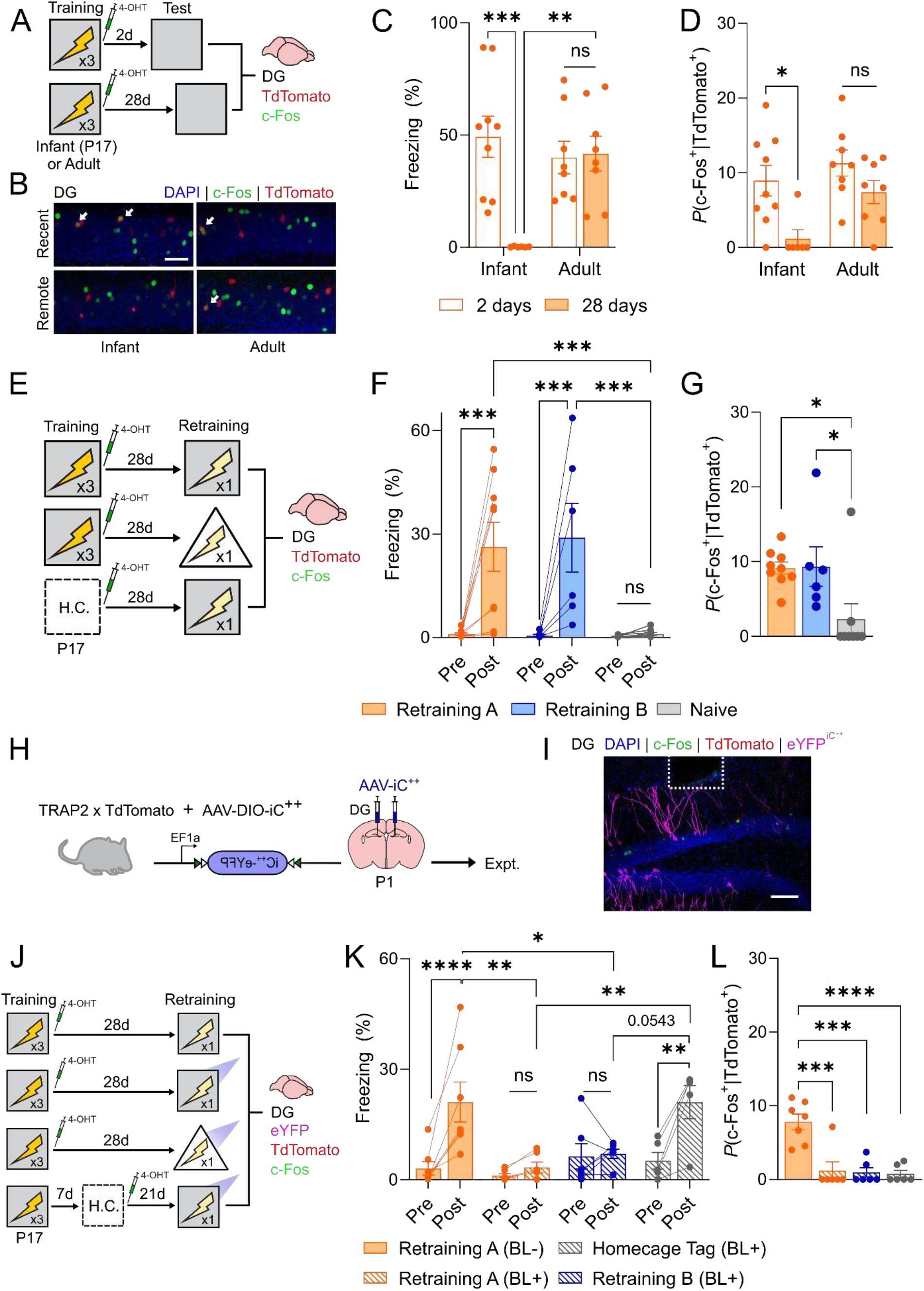
Infant learning produces latent engrams that facilitate subsequent learning during adulthood. (A) Infant (P17) or adult (P56-70) TRAP2xTdTomato mice were trained using contextual fear conditioning and tagged with 4-OHT, tested either 2 or 28 days later, and perfused after 90 minutes post-testing. (B) Representative images showing training-tagged (TdTomato^+^; red) and test activated (cFos^+^; green) neurons in the dentate gyrus. Arrows indicate engram neurons that were reactivated during testing (i.e., c-Fos^+^TdTomato^+^). Scale bar, 50 μm. (C) Infant-trained mice froze more at a recent delay than a remote delay (two-way ANOVA, main age effect: *F*_1, 27_ = 4.34, *p* = 0.0468; main delay effect: *F*_1, 27_ = 9.28, *p* = 0.0051; age x delay interaction *F*_1, 27_ = 10.74, *p* = 0.0029). (D) Infant-trained mice show increased reactivation of training-tagged dentate gyrus cells at a recent delay compared to a remote delay (two-way ANOVA, main age effect: *F*_1, 27_ = 5.81, *p* = 0.0230; main delay effect: *F*_1, 27_ = 10.72, *p* = 0.0029; age x delay interaction *F*_1, 27_ = 1.19, *p* = 0.2855). (E) Trained or naive (i.e., home cage; H.C.) infant TRAP2xTdTomato mice were tagged with 4-OHT, then re-trained 28 days in context A or B, and perfused after 90 minutes. (F) Infant-trained mice froze more than naive mice following re-training in either the A or B contexts (two-way ANOVA, main re-training effect: *F*_1, 20_ = 23.40, *p* = 0.0001; main group effect: *F*_2, 20_ = 5.48, *p* = 0.0127; re-training x group interaction *F*_2, 20_ = 5.52, *p* = 0.0123). (G) Infanttrained mice show increased reactivation of training-tagged dentate gyrus cells following retraining in contexts A or B (one-way ANOVA, main group effect: *F*_2, 20_ = 5.01, *p* = 0.0173). (H) Mice were microinjected with AAV-IC^++^ in the dentate gyrus at P1. (I) Representative images showing AAV-IC^++^ expression in the dentate gyrus with optic fibers implanted above. Scale bar, 100 μm. (J) Infant TRAP2xTdTomato mice were trained and tagged with 4-OHT, then re-trained 28 days later with either no light (control) or blue light to silence dentate gyrus training-tagged neurons and perfused after 90 minutes. (K) Infant-trained mice froze less following re-training across contexts when training-tagged dentate gyrus cells were inhibited compared to homecage-tagged or non-inhibited controls (two-way ANOVA, main re-training effect: *F*_1, 20_ = 28.79, *p* < 0.0001; main group effect: *F*_3, 20_ = 3.58, *p* = 0.0322; re-training x group interaction *F*_3, 20_ = 7.18, *p* = 0.0019). (L) Infant-trained mice show decreased reactivation of infant training-tagged dentate gyrus cells following dentate gyrus opto-inhibition (one-way ANOVA, main group effect: *F*_3, 21_ = 15.22, *p* < 0.0001).

Mirroring this behavioral pattern, engram reactivation (*P*[c-Fos|TdTomato]) at the remote test was reduced in mice trained as infants but not in mice trained as adults (Fig. 1D and Fig. S1A and B).

Together, these findings indicate that contextual fear memories formed in infancy are not accessible at remote timepoints and show little evidence of engram reactivation. However, retrieval failure does not necessarily imply memory loss. A large body of work in both humans and animals suggests that experiences that are not accessible to explicit recall can nonetheless persist in a latent form and influence subsequent learning^5^. We therefore asked whether dentate gyrus engrams supporting infant-acquired memories are reengaged during a later retraining phase, potentially facilitating performance by providing a scaffold for relearning and related learning.

TRAP2 × TdTomato mice were trained in contextual fear conditioning and treated with 4-OHT immediately after training to permanently label ensembles activated during encoding. One month later, mice were retrained either in the original context (context A) or a similar context (context B) to assess facilitation of relearning and related learning, respectively. During this retraining phase, a single shock was delivered after 150 s, and freezing was measured before (Pre) and after (Post) shock delivery. Control mice were treated identically but did not undergo training at P17 (ensemble labeling occurred in the home cage; Fig. 1E).

Mice exhibited minimal freezing prior to shock delivery during the retraining phase, confirming that infant-acquired fear memories were not retrieved under these conditions (i.e., infantile amnesia).

Despite this, shock presentation elicited post-shock freezing selectively in mice with prior infant training (Fig. 1F). Retraining induced memory (Fig. S2A and B) and facilitated relearning was observed at different training-retraining delays (Fig. S2C and D) and across developmental ages (Fig. SE and F). Because postshock freezing reflects a conditioned rather than unconditioned response to shock^7^, this effect is consistent with the idea that infant-acquired memories facilitate subsequent learning in adulthood. Using activitydependent engram tagging, we found that this facilitation was accompanied by reactivation of infantacquired dentate gyrus engrams, as indexed by increased overlap between TdTomato+ and c-Fos+ populations (Fig. 1G; Fig. S1C and D).

Notably, this facilitation was observed in both the original (context A) and similar (context B) environments, indicating that prior infant experience enhances learning even in related contexts. Facilitation in context B may suggest that infant-acquired contextual representations are relatively imprecise^8^, consistent with the idea that they form schema-like representations that generalize across contexts. In line with this, in separate cohorts we found that infant-acquired context memories are less precise than those formed in adulthood; following conditioning in context A at P17, mice exhibit equivalent freezing in contexts A and B when tested 48 hours later (Fig. S2G-I).

To test whether infant-acquired dentate gyrus engrams were required for facilitation, we selectively inactivated them during adult learning. An adeno-associated virus (AAV) expressing the inhibitory opsin (iC^++^) was microinjected into the dentate gyrus of TRAP2 mice. Administration of 4-OHT immediately after training permanently tagged ensembles activated during encoding with this blue-light inhibitory opsin (Fig. 1H-J). Subsequent BL inhibition of these ensembles (and not a control ‘home cage’ neuronal ensemble) prevented reactivation of the infant-acquired engram (Fig. 1K; Fig. S1E and F) and attenuated post-shock freezing during retraining in either the original context (context A) or a similar context (context B) (Fig. 1L).

Our findings highlight a central paradox of infantile amnesia: memories formed early in life become inaccessible yet go onto influence related learning in adulthood. Using activity-dependent engram tagging and causal manipulations, we show that early-life dentate gyrus engrams—though behaviorally silent— persist as functional substrates that can scaffold later learning if reactivated by related experiences during adulthood. Consistent with this view, similar evidence for infant-acquired memories supporting relearning and related learning has been reported in mice^9^ and rats^10^ using contextual and object-based paradigms. In particular, Bessières et al.^9^ reported closely related findings using comparable engram tagging and manipulation approaches. Our work was conducted independently and converges on the same conclusion.

However, where our findings differ from these previous studies is in terms of the locus of the schema-like substrate. Contreras et al^10^ and Bessières et al^9^ emphasized a time-limited role for the hippocampus in supporting relearning and related learning, instead highlighting prefrontal cortical contributions. In contrast, our results demonstrate a sustained role for the hippocampus, with activation of infant-acquired dentate gyrus engrams facilitating learning in both the original and related contexts up to one month following infant experience. Our results, along with previous studies^8,9^, identify an adaptive function for infantile memory formation and subsequent amnesia, which is to rapidly generate a neural scaffold in the brain that can serve as a foundation for subsequent experiences accumulated throughout life.

## Acknowledgements

This project was funded by an NSERC grant to P.W.F. C.M. was supported by a Restracomp award. We thank A. Ramsaran for comments on this manuscript.

## Author Contributions

Conceptualization, S.H., C.M. and P.W.F.; Methodology, S.H., C.M., S.A.J. and P.W.F.; Investigation, S.H. and C.M; Formal analysis S.H. and C.M.; Writing – Original Draft, S.H., C.M. and P.W.F.; Writing – Review & Editing, S.H., C.M., S.A.J., and P.W.F.; Funding acquisition – P.W.F.; Supervision, S.A.J., and P.W.F.

