## Supplemental Information for "Latent dentate gyrus engrams formed in infancy scaffold adult learning"

### Supplemental Experimental Procedures

All procedures were conducted in accordance with policies of the Hospital for Sick Children Animal Care Committee and conformed to both Canadian Council on Animal Care (CCAC) and National Institutes of Health (NIH) Guidelines on Care and Use of Laboratory.

#### Experimental animals

In these experiments, we used homozygous TRAP2 mice (Fostm2.1(icre/ERT2)Luo; C57BL/6N; acquired from Jackson Laboratory RRID: IMSR_JAX:030323) crossed with TdTomato reporter homozygous mice (Gt(ROSA)26Sortm14(CAG-tdTomato)Hze; C57BL/6N; acquired from Jackson Laboratory RRID:IMSR_JAX:007914) to produce F1 double heterozygous mice TRAP2::tdTomato. Mouse genotypes were determined by PCR analysis of tail DNA samples.

Mice were maintained on a 12-hour light/12-hour dark cycle at all times with lights on between 8 AM to 8 PM each day. Food and water were provided ad libitum. The litters, usually containing 4-12 pups, were weaned 21 days post date of birth and housed 2-5 mice per cage. Males and females were housed separately and were used in all experiments. “Infant” mice were 17 days old (i.e., P17) and “adult” mice were eight to ten-week-old (i.e., P56-70) at the initiation of  behavioural experiments, which occurred during the light phase.

#### Method details

#### Contextual fear conditioning apparatus

For the training, re-training, and testing, the conditioning context (“A”, gray square) had clear acrylic walls on the front and back of the chamber and aluminum walls on the sides and top of the chamber (32 x 25 x 25 cm). The floor consisted of metal grids that were used for administering foot shocks (0.5 mA, 2 s duration). A camera was installed above the chamber for monitoring behavior. The similar context (“B”, white triangle) had white acrylic walls inserted within the original conditioning context to create a triangular chamber (28 cm x 24 cm x 25 cm). The floor consisted of the metal grids that were used for administering foot shocks (0.5 mA, 2 s duration). Peppermint was applied to provide a distinct scent cue.

**Contextual fear conditioning procedures**

**Conditioning**

On the day of the conditioning, mice were taken from the vivarium and temporarily held in a quiet holding area for 60 mins. They were transferred to the conditioning room and placed into the conditioning chamber (context A). Shocks were delivered at 120 s, 150 s, and 180 s after placement in the context and mice remained in the chamber for an extra 60 s after the final shock. They were returned to the holding room for 60 mins, and then transferred back to the vivarium.

**Context tagging**

Immediately upon removal from the fear conditioning chambers, ensemble tagging was induced in mice via an intraperitoneal (i.p.) injection of 4-Hydroxytamoxifen (4-OHT; 25 mg/kg or 50 mg/kg injected at 5 µl/g mouse, for infants and adults, respectively). In “homecage” controls, mice were treated with 4-OHTin their home cage.

4-OHT powder was first mixed with 100% ethanol (0.25mg/6.25 µl or 1.5 mg/3.75 µl for infants and adults, respectively) and vortexed vigorously. The solution was then poured into a 50°C chamber, vortexing every 12 min until fully dissolved. An equal part cremaphore was added to create a stock solution that was stored at −20°C until required. On experimental days, the stock solution was mixed at a 1:4 or 1:2 ratio (for infants and adults, respectively) with PBS.

**Re-training and testing**

Mice were brought to the holding room 60 mins prior to re-training or testing. For the re-training conditions, a single shock was delivered 150 s after placement in the context and mice remained in the chamber for 150 s after the shock. In all conditions, mouse behavior was monitored via an overhead camera (3.75 frames/s), and freezing was automatically tracked using FreezeFrame software. For the re-training conditions, the percentage of time spent freezing for the first 150 s (i.e., pre-shock) and the last 150 s (i.e., post-shock) was used for statistical analyses. For all other conditions, the percentage of time spent freezing across the entire testing period was used for statistical analyses.

**Optogenetic engram inhibition**

Mice were connected to optic cables and placed within a neutral homecage for 5 mins the day before the session to habituate to the cables. During the re-training session, photoinhibition (473 nm, 7-10 mW) was delivered for 30 s starting at 135 s (15 s before shock administration), and freezing was scored manually (because the presence of optical cables precluded automated scoring). Mice in the no light conditions were tested identically, except that light stimulation was not delivered.

**Viral vectors**

**AAV(DJ)-EF1a-DIO-iC$^{++}$-EYFP (AAV-IC$^{++}$).** The plasmid was kindly provided by Dr. Karl Dessieroth (Stanford University) and was packaged in-house. The virus was used to tag and optogenetically silence engram neurons in DG in TRAP2 mice. iC$^{++}$, a blue-light sensitive inhibitory opsin, is inverted and flanked by two Lox sequences, which prevents expression. However, in the presence of active Cre recombinase from the TRAP2 system, the Lox sequences are recognized and the iC$^{++}$ sequence is reverted to the coding status leading to expression of the transgene. An EYFP (enhanced yellow fluorescent protein) allows visualization of the opsin. The expression of the vector is driven by EF1α (human elongation factor 1- subunit α) promoter which is a neuron-specific promoter and can elicit expression in all neuronal types ubiquitously.

**Stereotaxic surgery**

**Virus microinfusion**

Neonatal virus injection procedures were adapted from previous studies[1]. P1 mice were anesthetized through hypothermia and mounted to a chilled metal plate using laboratory tape. The scalp was incised and retracted, and the glass micropipette connected to a nanoliter injector (Nanoject III, Drummond Scientific) was used to pierce the skull above dorsal DG (approximate coordinates AP +1.0 mm, ML ±0.8, DV -1.35 mm from lambda). The pipette injected 0.12-0.15 $\mu$l AAV over 1 min and remained in place for an additional 3 min. The scalp was sealed with Vetbond Tissue Adhesive (3M) and covered with Polysporin. Pups remained on the heating pad until all surgeries were completed and continuously monitored until mobile, at which time the litter was returned to the dam.

**Optrode Placement**

Before surgery, mice were treated with atropine sulfate (0.1 mg/kg) and chloral hydrate (400 mg/kg) via intraperitoneal (i.p.) injection. Once a stable level of anesthesia occurred, mice received a subcutaneous (s.c.) injection of Meloxicam (4 mg/kg) for post-surgery analgesia. The top of the scalp was disinfected with ethanol (70%), the scalp incised and retracted. A hole was made above the region of interest using an electric hand drill and an optical implant was inserted into the region of interest. To target the dentate gyrus the following coordinates were used (with respect to bregma): AP -2.2 mm, ML ±1.5 mm, DV -2.3 mm. The optical implant was constructed by attaching a fiber (200 μm diameter, 0.37 numerical aperture, Thorlabs) to a ceramic zirconia ferrule (1.25 mm diameter, Thorlabs) with epoxy resin. Two fibers were implanted bilaterally above injection sites on either side of the brain. Implants were secured on the skull by the application of black dental cement and screws. The wound was treated with Polysporin and the mice received a 1 ml injection of saline (0.9%, s.c.) for hydration. During recovery, mice were placed on a clean surface with access to a heating pad. After recovery, the mice were transferred to a clean cage and returned to the vivarium. The health of the mice was monitored for 3 days post-surgery. One day after surgery, mice were treated with an additional subcutaneous (s.c.) injection of Meloxicam (4 mg/kg) for post-surgery analgesia. The behavioral procedures took place approximately 1 week after surgery.

**Perfusion, tissue processing and confocal imaging**

**Perfusions**

At 90 min following the completion of behavioural testing, mice were perfused transcardially with 40 ml of cold phosphate buffered saline (PBS; 0.1 M) followed by 40 ml of cold paraformaldehyde (4%; PFA). PBS and PFA volumes were adjusted for different mouse ages. Brains were extracted, post-fixed in PFA overnight and stored in PBS with 0.3% sodium azide. Brains were switched to an anti-freezing solution (30% sucrose solution) until they submerged, and sliced coronally in 50 µm-thick sections in a cryostat (Leica CM1850). The sectioned tissues were transferred to PBS (0.1 M) in well plates and stored for further processing. In the case of long-term storage, the solution was exchanged with a PBS solution that contained NaN3 (0.2%). All tissues were stored at 4 °C.

**Immunohistochemistry**

For experiments involving quantification of the number of cells positive for immunofluorescence or immunofluorescence signal intensity, all staining was performed at once using the same antibody solutions. Free-floating sections were blocked with PBS containing 4% normal goat serum and 0.5% Triton-X for 1 h at room temperature. Afterwards, sections were incubated with primary antibodies in fresh blocking solution: rabbit anti-c-Fos (1:1000, Synaptic Systems, cat# 226003), chicken anti-GFP (1:1000, Aves, cat#

GFP-1010) for 24 hours (for c-Fos experiments) at 4 °C. Sections were washed three times for 15 min each with PBS containing 0.1% Tween-20 (PBST), then incubated with PBST containing secondary antibodies: goat anti-chicken Alexa Fluor 488 (1:500, Invitrogen, cat# A-11039, goat anti-rabbit Alexa Fluor 488 (1:500, Invitrogen, cat# A-11008), goat anti-rabbit Alexa Fluor 568 (1:500, Invitrogen, cat# A-11011) for 2 h at room temperature. Sections were washed with PBS, counterstained with DAPI, mounted on gel-coated slides, and coverslipped with Permafluor mounting medium (ThermoFisher Scientific, cat# TA-030-FM).

**Confocal microscopy**

We obtained all images using a confocal laser scanning microscope (LSM 710; Zeiss). We acquired images by scanning 20 z-stacks (1 μm steps) per section with a 20X objective and tiling to cover the region of interest. The laser parameters (e.g. laser power, pinhole size, multiplier gain, and any other digital parameters) were kept the same for all the acquired images of each experiment. The imaging parameters were set using a sample from the control group. The location of each image was selected according to markers of interest per mouse brain atlas.

**Quantification and statistical analyses**

**Cell counting**

For cell counts, the number of cells expressing the marker of interest was quantified. Marker-positive cells in the entire cross-section of the region of interest were counted. All images underwent identical thresholding using the same parameters for each experiment. First, the z-stacks were superimposed, background noise filtered and signals thresholded to acquire countable masks. All the processing was done in Fiji software (National Institute of Health). The counts from the replicate images from each animal were combined to create a final average for each mouse.

For each mouse, cells were counted manually in Fiji (National Institutes of Health) using 5 images acquired from 3-5 sections and averaged. To estimate the number of $DAPI^+$ cells in the DG granule layer, $DAPI^+$ cells were counted in a small volume (approximately 10 × 50 × 15 $\mu$m3) to obtain the $DAPI^+$ density ($mm^{-3}$). For all experiments, the volume of the granule layer within each image was measured and multiplied by the $DAPI^+$ density value for the appropriate age group to obtain the estimated $DAPI^+$ cell number. Quantification of c-$Fos^+$ and $TdTomato^+$ cells was limited to the granule layer. The proportion of granule layer cells expressing TdTomato or c-Fos after contextual fear conditioning training, re-training, test, or in the home cage are reported as $P(TdTomato^+|DAPI^+)$ and $P(c\text{-}Fos^+|DAPI^+)$, respectively. Engram reactivation rates were computed as $P(TdTomato^+c\text{-}Fos^+|TdTomato^+)$.

**Statistical analysis**

The sample size for the experimental groups was determined based on previous publications[1-3]. Mice were pseudo-randomly assigned to different groups with roughly equal numbers of male and female mice. The experimenter was blinded to group IDs when quantifying images and behavioral outcomes. Cell counts were normalized to either DAPI or TRAP counts. The DAPI normalization was used when the overall population was the denominator of interest, and the TRAP normalization was used when the engram population was the denominator of interest. In all experiments, data was analyzed using either one-way or two-way analysis of variance (ANOVA), with repeated measures followed by post-hoc tests (Sidak or Tukey) when appropriate. Statistical significance was set to 0.05. Different mice were used for all experiments. Statistical analysis was carried out using GraphPad Prism (version 8.0.1).

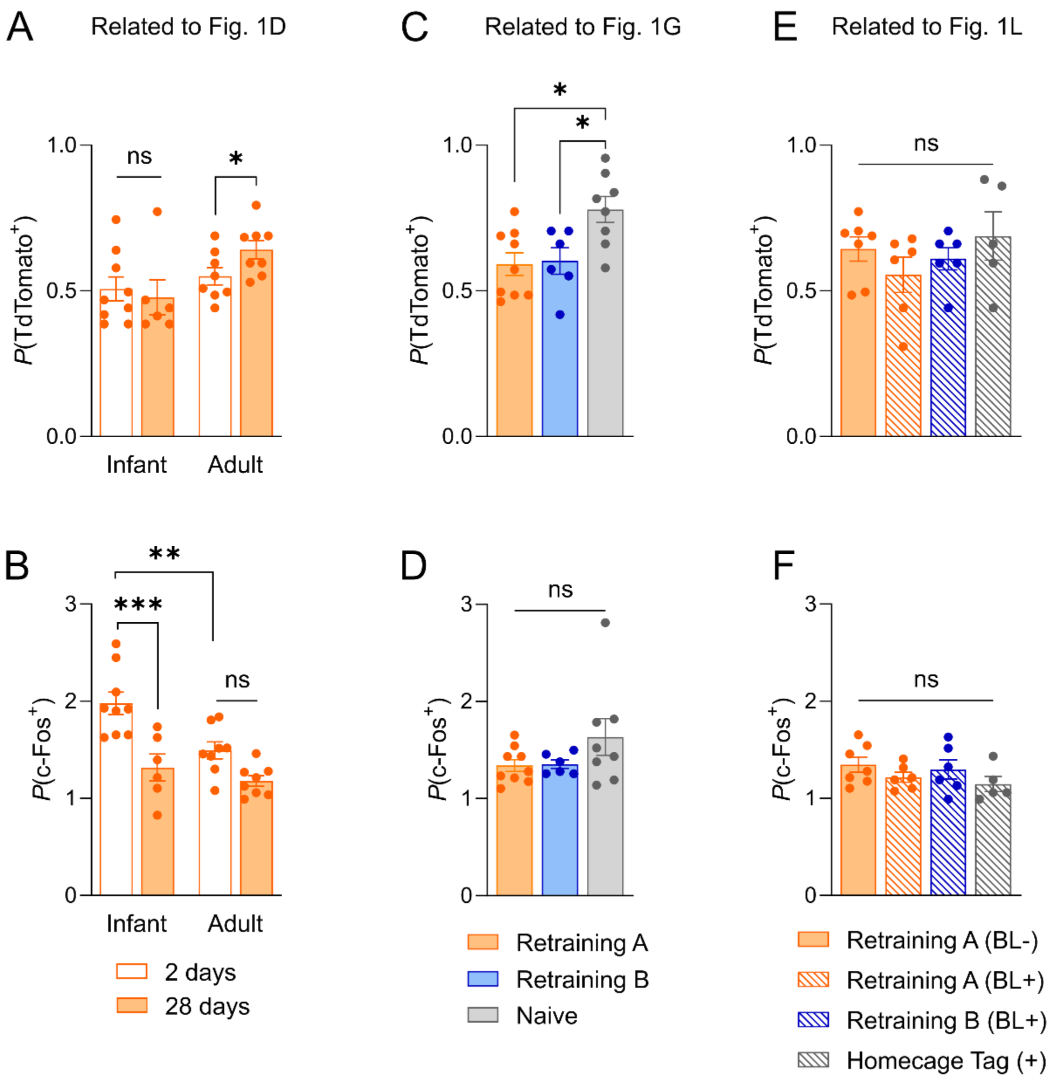


**Supplemental Figure 1. Training (TdTomato⁺) and retreaining (c-Fos⁺) quantification.**
(A) Adult mice in the remote testing condition show an increased proportion of training-

tagged dentate gyrus cells (two-way ANOVA, main age effect: $F_{1, 27} = 6.53$, $p = 0.0166$; main delay effect: $F_{1, 27} = 0.59$, $p = 0.4502$; age x delay interaction $F_{1, 27} = 2.19$, $p = 0.1509$). (B) Infant mice in the recent testing condition show increased proportion of c-Fos$^{+}$ cells in the dentate gyrus (two-way ANOVA, main age effect: $F_{1, 27} = 9.35$, $p = 0.0050$; main delay effect: $F_{1, 27} = 22.92$, $p < 0.0001$; age x delay interaction $F_{1, 27} = 1.19$, $p = 0.0993$). (C) Naive mice show an increased proportion of training-tagged dentate gyrus cells (one-way ANOVA, main group effect: $F_{2, 20} = 6.26$, $p = 0.0078$). (D) All groups show a similar proportion of c-Fos$^{+}$ cells in the dentate gyrus (one-way ANOVA, main group effect: $F_{2, 20} = 1.87$, $p = 0.1805$). (E) All groups show a similar proportion of training-tagged dentate gyrus cells (one-way ANOVA, main group effect: $F_{3, 20} = 0.97$, $p = 0.4257$). (F) All groups show a similar proportion of c-Fos$^{+}$ cells in the dentate gyrus (one-way ANOVA, main group effect: $F_{3, 20} = 1.20$, $p = 0.3349$).

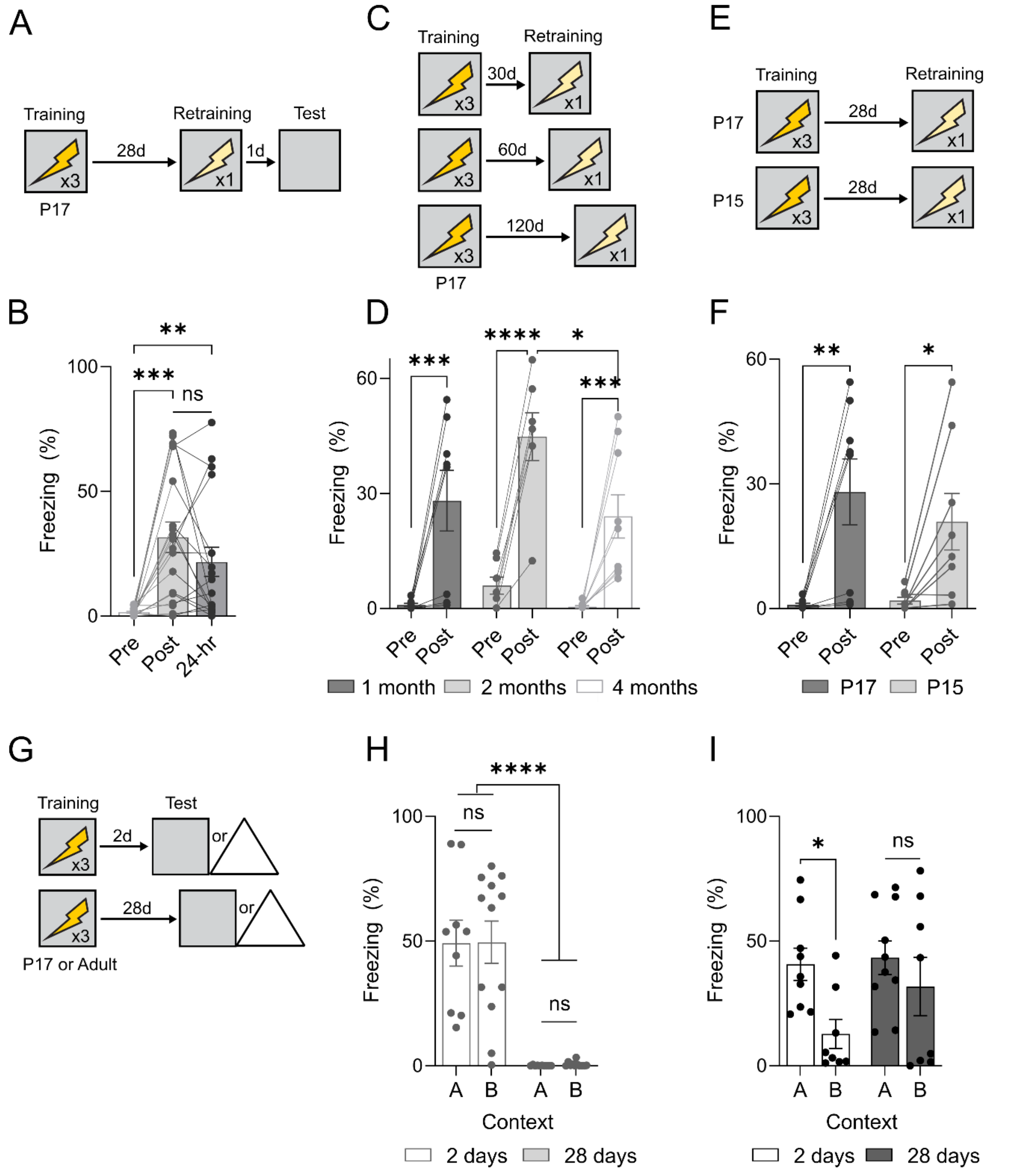


**Supplemental Figure 2. Infant learning produces persistent, generalizable memories.** (A) Infant TRAP2xTdTomato mice were trained, re-trained after 28 days, and tested 1 day later. (B) Infant-trained mice froze more immediately following re-training and during test the next day (one-way ANOVA, main session effect: $F_{1.987, 33.78}$ = 13.04, $p$ < 0.0001; main subject effect: $F_{17, 34}$ = 2.006, $p$ = 0.0412). (C) Infant TRAP2xTdTomato mice were trained and re-trained either 30, 60, or 120 days later. (D) Infant-trained mice froze more following re-training across all delays (two-way ANOVA, main re-training effect: $F_{1, 21}$ = 68.68, $p$ < 0.0001; main group effect: $F_{2, 21}$ = 3.55, $p$ = 0.0471; re-training x group interaction $F_{2, 21}$ = 1.57, $p$ = 0.2312). (E) Infant TRAP2xTdTomato mice were trained at either P17 or P15 and re-trained 28 days later. (F) P17 and P15 infant-trained mice froze more following re-training (two-way ANOVA, main re-training effect: $F_{1, 20}$ = 27.41, $p$ < 0.0001; main group effect: $F_{2, 20}$ = 0.20, $p$ = 0.8212; re-training x group interaction $F_{2, 20}$ = 0.34, $p$ = 0.7151). (G) Infant (P17) or adult (P56-P70) TRAP2xTdTomato mice were trained and tested 2 or 28 days later in Context A or B. (H) Infant-trained mice froze more at the 2 day delay, with no difference across contexts (two-way ANOVA, main context effect: $F_{1, 38}$ = 0.01, $p$ = 0.9447; main delay effect: $F_{1, 38}$ = 61.45, $p$ < 0.0001; context x delay interaction $F_{1, 38}$ < 0.01, $p$ = 0.9907). (I) Adult-trained mice froze more in context A than B after a 2 day delay, but not a 28 day delay (two-way ANOVA, main context effect: $F_{1, 31}$ = 6.33, $p$ = 0.0173; main delay effect: $F_{1, 31}$ = 1.88, $p$ = 0.1798; context x delay interaction $F_{1, 31}$ = 1.08, $p$ = 0.3059).
